# Head motion correction shapes functional network estimates: evidence from healthy and Parkinson’s disease cohorts

**DOI:** 10.1101/2022.12.26.520413

**Authors:** Francesca Saviola, Stefano Tambalo, Donna Gift-Cabalo, Lisa Novello, Enrica Pierotti, Giuseppe Rabini, Alessandra Dodich, Luca Turella, Dimitri Van De Ville, Jorge Jovicich

## Abstract

An open discussion in studies of intrinsic brain functional connectivity is the mitigation of head motion-related artifacts, particularly in the presence of peculiar symptomatology such as in Parkinson’s disease (PD). Previous studies show that Independent Component Analysis (ICA) denoising improves the reproducibility of functional connectivity findings by detecting sources of non-neural signals. However, there is still no consensus about which pre-processing pipeline should be applied in natural high motion populations such as PD, particularly in relation to novel functional network descriptions derived from dynamic connectivity analyses. In this study, we investigated how different pre-processing pipelines affect intrinsic brain connectivity metrics, both static and dynamic, derived from a group of young healthy controls (HC) and a group of PD participants.

A total of 20 HC and 20 PD subjects participated in this 3 T MRI study. Resting-state functional MRI images were used to test the effects of the pre-processing pipeline of static (sFC) and temporal-varying functional connectivity (dFC) estimations. Both MRI datasets were pre-processed using three different workflows differing in the motion correction approach: (i) standard motion realignment (*mc*); (ii) motion outlier detection and deweighting based on image intensity change estimations (*DVARS*) and (iii) ICA-based noise removal using reference noise features (*AROMA*). Furthermore, the PD dataset was also processed with a fourth method by applying an ICA-based denoising (*FIX*), previously trained on the HC group. sFC analysis was performed using Group ICA, by temporally concatenating different pre-processing types in pairs of different runs. Two types of dFC analyses were considered: innovation-driven co-activation patterns (iCAPs) and co-activation patterns (CAPs). CAPs allow dFC estimations that do not require the deconvolution of the hemodynamic response function and its derivative, thus potentially being less sensitive to head-motion related noise. We found that regardless of substantial head motion differences in the two groups, sFC results were consistent across denoising strategies. Conversely, dFC was extremely sensitive to denoising strategies, particularly for the PD group with the transient-based dFC analyses. Indeed, the use of the peak-based dFC framework enables the detection of time-varying networks but in a way that is highly dependent on the motion correction pipeline. In conclusion, we show that dynamic functional network representations are highly sensitive to both head motion and to fMRI denoising methods. These findings stress the importance of considering and reporting these experimental aspects to help with the reproducibility and interpretation of different studies. Future work is needed to further investigate transient-based dFC strategies that are more robust to head motion.

## 1. Introduction

### 1.1 Background

Head motion is one of the majors confounds in brain functional MRI (fMRI) studies, particularly in studies of functional connectivity (FC) networks derived from intrinsic activity ^1–3^. The impact of head motion, however, has until now been mostly investigated considering static brain networks, which are representations of FC that use the entire timecourse of the fMRI acquisition ^4^. Little is known about how head motion affects dynamic FC network representations, especially in the case of frame-wise techniques.

Some of the first studies in the field highlighted that even small or transitory head movements could result in severe biases in static FC estimations ^5,6^ and consequently misleading interpretations in the context of pathologies ^7,8^. However, while in the case of static FC the need to carefully correct for motion is widely recognized across experts, in the context of dynamic FC this is still a matter of debate. On one hand, some studies suggest that head movement may only partially affect dynamic metrics and their reliability ^9^, whereas others claim the need of a specific tool to effectively correct for motion and overcome biases in the dynamic FC estimates ^10^. Overall, there is a strong interest in the fMRI neuroimaging community to develop robust processing strategies aimed at controlling head motion confounds in the BOLD time course ^11^, but an understanding of their impact on dynamic FC networks is still lacking.

At the current state-of-the-art, several methodologies have been proposed as denoising pipelines for BOLD fMRI time series. These pipelines are typically set up as a “cascade” of various possible steps for which, to the best of our knowledge, there is no global consensus. The first and most common way of correcting for head motion is the so-called linear realignment ^12^ retrospectively done after the acquisition (here referred to as <u>motion correction pipeline; *mc*</u>). In this traditional method, head motion estimates over time are calculated along three translational trajectories (left-right, anterior-posterior and dorsal-ventral) and three rotational planes (roll, pitch and yaw). Following the *mc* correction step, data can be further corrected by discarding or weighting the BOLD signal of time points based on their level of head motion level. Common head motion metrics used for this include: (i) framewise displacement^5^ (FD), defined as the average frame-to-frame whole-brain displacement considering both rotation and translation parameters differences between consecutive frames; (ii) DVARS ^5^, defined as the root mean square (RMS) intensity difference of single volume (N) to another single volume (N+1) (here referred to as <u>motion outliers deweighting pipeline; *DVARS*</u>); (iii) RMS intensity difference of single-volume respect to the reference volume. Subsequently, brain fMRI volumes defined as corrupted by the chosen head motion metric may be fed into a confound matrix and regressed out from the BOLD time series with a general linear model. A weakness of the above-cited metrics is the implicit assumption that motion remains similar along the whole fMRI time course, with the extreme case of framewise displacement, which is considered to be stationary even across translational and rotational planes thus neglecting its spatio-temporal features ^13,14^.

In an attempt to address these limitations, especially in the case of populations that tend to present large head motion (such as pediatrics or clinical cohorts), distinct denoising pipelines based on Independent Component Analysis (ICA) have been proposed, including ICA-AROMA ^15^ and FMRIB’s ICA-based Xnoisifier (FIX; Salimi-Khorshidi et al., 2014)^16^. In contrast to previous methods, ICA-derived denoising pipelines are completely data-driven, can identify different sources of structured noise, and unlike brain volume censoring, they preserve the temporal structure of the data. Specifically, ICA-AROMA (here referred to as <u>motion denoising; *AROMA*</u>) is an algorithm used for the automatic removal of motion-related components based on their spatial and temporal features. Conversely, FIX (here referred to as <u>motion denoising;</u> *<u>FIX</u>*) is an fMRI noise detection algorithm that uses binarized characterization of *‘good’* or *‘bad’* independent components (IC) based on ∼180 features captured from IC’s spatial and temporal characteristics through a multi-level classifier^17,18^.

In sum, when interested in studying functional brain networks, there is a wide variety of different and publicly available head motion correction strategies that can be applied. However, it is not clear how these approaches may differ between populations that show different levels of natural head motion (natural motion intended as not the result from motion instructions to healthy collaborative subjects).

In this study, we investigate how intrinsic functional networks derived from both static and dynamic FC fMRI methods are affected by head motion. To this end, we manipulated denoising strategies on two populations with different natural head motion levels: healthy young volunteers (where we tested *mc, DVARS, AROMA*) and Parkinson’s disease participants (where we tested *mc, DVARS, AROMA* and *FIX* trained on the healthy volunteers).

## 2. Materials and Methods

### 2.1 Participants

A total of 20 healthy controls (HC - 10 females; age 24±3 years) and 20 PD subjects (8 females; age 67±7 years) participated in this study, which was approved by the Ethical Committee of the University of Trento, Italy. For what concerns HC, exclusion criteria included the presence of any neurological and/or psychiatric disease. For the PD participants inclusions criteria were diagnosis of idiopathic PD based on the Movement Disorders Society-Unified Parkinson disease rating Scale (MDS) criteria ^19^, with disease severity < 3 based on the modified Hoen and Yahr scale ^20^ and being under anti-parkinsonian medication. Patients with evidence of dementia or other neuropsychiatric disorders were excluded. All patients were tested while in their medication-on condition. The participants from the two cohorts gave written informed consent.

### 2.2 MRI Acquisition

Data were acquired with a 3T clinical MRI scanner (MAGNETOM Prisma, Siemens Healthcare, Erlangen, Germany) equipped with a 64-channel receive-only head-neck RF coil. Structural T1-weighted multi-echo MPRAGE ^21^ for the PD cohort (TR=2.5s, TEs=[1.69, 3.55, 5.41, 7.27 ms], 1mm-isotropic voxels), standard MPRAGE for the HC group (TR/TE=2.31s/3.48ms, 1mm-isotropic voxels) and resting-state functional MRI (rs-fMRI, TR/TE=1s/28ms, 3mm-isotropic voxels, FA=59 degree, MB=6) images were used to test the effects of the motion-denoising approach on static (sFC) and time-varying FC (dFC) estimations. Double-echo gradient echo sequence was acquired for both cohorts to later control con geometric distortions (TR 682ms, TE1/TE2 4.2/7.4ms, 3mm-isotropic voxels).

### 2.3 MRI Pre-processing

The MRI datasets from the two groups (HC and PD) were pre-processed using three different workflows (see Figure 1). Common pre-processing steps across the three workflows were conducted with an FSL-based automated pipeline (https://github.com/tambalostefano/lnifmri_prep) and included: (1) slice timing and head motion correction (*mc*); (2) co-registration of the T1-weighted image to the rs-fMRI time-series; (3) T1-weighted image tissue segmentation; (4) rs-fMRI temporal detrending and median filtering; (5) regression from the rs-fMRI time-series of the 6 head motion parameters, white matter and cerebro-spinal fluid signals (*<u>standard motion correction, mc</u>*); (7) normalization to standard MNI template space; and (8) spatial smoothing 6 mm FWHM Gaussian kernel size.

**Figure 1:**
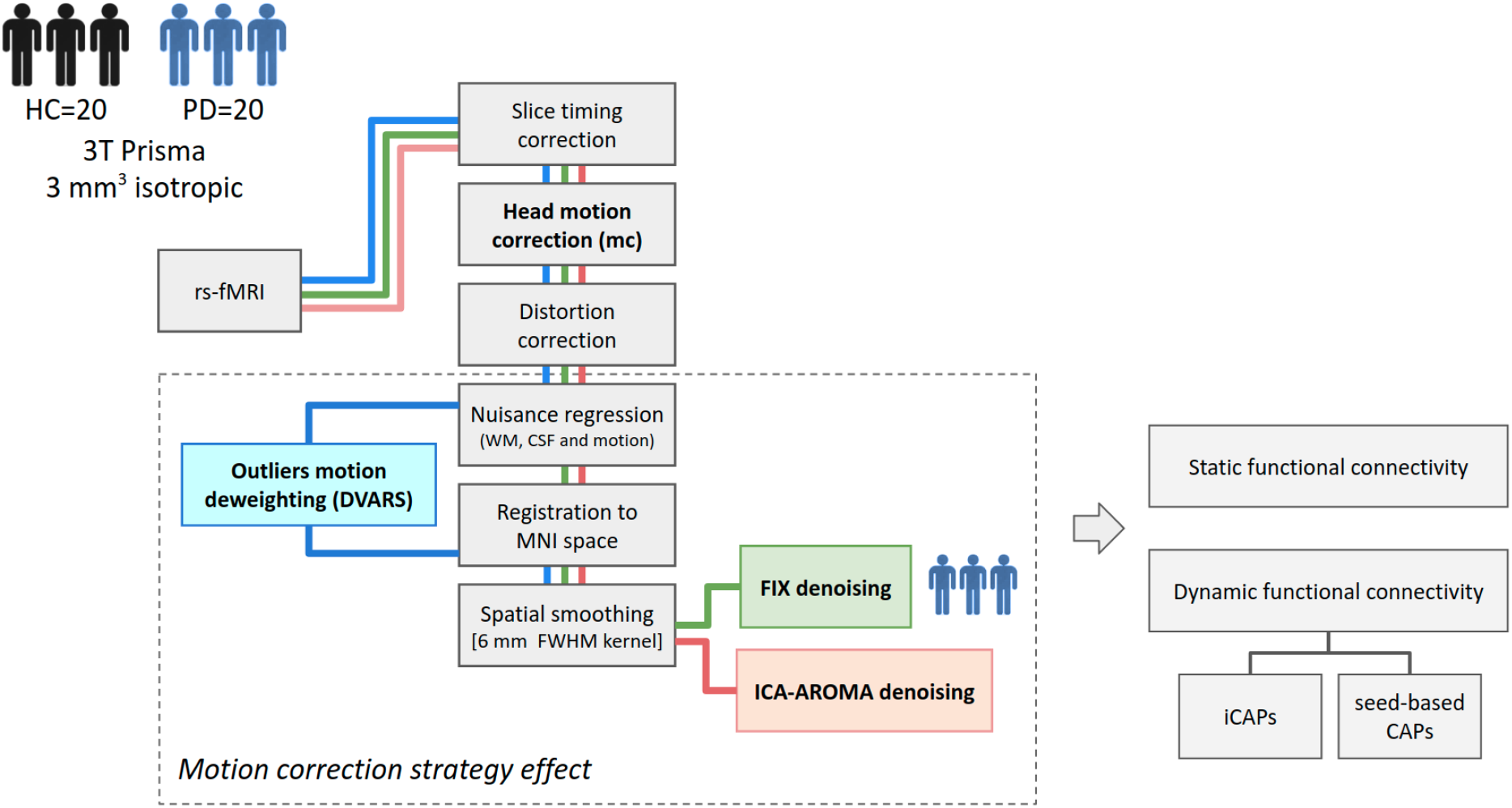
Schematic representation of the pre-processing pipelines for the two cohorts, healthy controls (HC) and Parkinson’s disease participants (PD). The pre-processing pipeline of resting-state fMRI (rs-fMRI) data for HC included three different workflows: (i) *<u>mc</u>*: head motion correction (in grey); (ii) *<u>DVARS</u>*: outliers’ motion deweighting (in light blue); (iii): *<u>AROMA</u>*: ICA-AROMA denoising (in light pink). For the PD sample, the same three pipelines were used together with another denoising technique: *<u>FIX</u>* (in light green).

Together with these standard pre-processing steps, the three pipelines differed in the following motion correction approaches: (i) *mc* (nothing additional to the standard pre-processing); (ii) motion outliers detection and deweighting based on DVARS estimations (*<u>DVARS</u>*; Power et al., 2012); (iii) noise removal by means of ICA-AROMA (*<u>AROMA</u>*), where motion-related artifacts are detected automatically if the high-frequency content exceeds 35% of the fMRI time course, Cerebro-spinal fluid fraction is > 10%, and if it falls behind the decision boundary created for edge fraction and maximum correlation with realignment parameters ^15^.

Additionally, the PD cohort was further pre-processed (iv) using FIX-Classifier (*<u>FIX</u>*), previously trained on 17 subjects of the HC dataset ^17,22^. The definition of a training set to train the classifier consisted in (1) ICA decomposition of rs-fMRI time series into n=30 independent components, (2) manual classification of motion-related components by visual inspection of their spatiotemporal features. The training set was then used to train the classifier and the accuracy was assessed by a leave-one-out approach. The *FIX* classifier was further applied to the PD dataset with a threshold of 20 corresponding to 95.2% (mean TPR) of correctly classified components.

### 2.4 Static functional connectivity

In order to estimate static FC differences across the different head motion pipelines, the pre-processed resting-state fMRI data was then decomposed using Multivariate Exploratory Linear Optimized Decomposition into Independent Components 3.0 (MELODIC) into their distinct spatial and temporal components using ICA. As the aim of the analysis was to detect resting-state FC changes associated with different head motion pre-processing pipelines, we did not assume consistent temporal responses within subjects. Therefore, the ICA group analysis was temporarily concatenated (FSL; Jenkinson, Beckmann, Behrens, Woolrich, & Smith, 2012) in different pre-processing pipeline runs, respectively combining *mc* vs *DVARS, mc* vs *AROMA* and *mc* vs *FIX* (for the PD only), to then perform a paired nonparametric permutation test in each group separately. The ICs number was manually set to 20 as low-order model analysis ^24^. To separate noise components from the underlying resting-state networks, ICs were tested for their correlation (threshold of r-value > 0.2) to labelled networks ^25^. Subsequently, 10 ICs out of 20 were retained with the highest r-value. As a final step in the network identification, ICs were visually inspected by expert users to detect consistency between ICs matching with template networks (high correlation values) resembling well-known functional networks.

### 2.5 Dynamic functional connectivity

Concerning the evaluation of head motion effects and denoising approaches on dynamic FC, the following two frame-wise methods were considered.

#### 2.5.1 Transient-based dynamic FC analysis

We first estimate recurrent brain states by means of innovation-driven co-activation patterns (iCAPs; Karahanoğlu & Van De Ville, 2015) (https://c4science.ch/source/iCAPs/). This technique enables the detection of iCAPs in a data-driven way, by looking at transients of the fMRI signal. Indeed, by computing at the single-subject level the derivative of the previously regularized and hemodynamic response function (HRF) deconvolved fMRI signal, the innovation signal is derived, comprehending the original fluctuations of the BOLD time course ^27^. These innovations signals are then clustered together across subjects, in order to reconstruct the iCAPs, which describe how brain regions are functionally characterized by similar temporal dynamics ^28^.

The iCAPs framework was applied separately to the two groups (HC and PD, respectively) for all the types of pre-processing and by means of consensus clustering. The best-fitting k was chosen for both populations in k=10. As a final step, spatiotemporal transient-informed regression ^29^ was used to reconstruct the time course of each iCAPs. Temporal properties of iCAPs for each group in each pre-processing type, were extracted by transforming the time courses to z-scores and thresholding them (|z-score|>1), and total duration of each iCAPs were retrieved (i.e. duration of the overall iCAP activation represented as percentage of the scanning time) ^30^. In order to separate noise components from the underlying dynamic networks, ICAPs were tested for their correlation (threshold of r-value > 0.2) to labelled networks ^25^

#### 2.5.2 Peak-based dynamic FC analysis

The results obtained by the transient-based dFC analysis, showed large effects of denoising methods on iCAPs estimation, especially in the PD cohort. We hypothesized that this may be due to the incorrect assumption of the HRF model when applied to a BOLD time series with strong frame-wise head motion effects. To test this, in the PD group we also applied a framewise dFC method that was not dependent on assumptions of the HRF for the estimation of brain recurrent states: co-activation patterns (CAPs; Liu & Duyn, 2013) (https://c4science.ch/source/CAP_Toolbox.git.). In our case, CAPs were estimated by defining a seed in the posterior cingulate cortex (PCC) region from the Harvard-Oxford Labeling atlas and extracting co-activations and co-deactivations patterns with respect to the seed, for limited periods of the time course, with temporal clustering aiming at detecting the Default-mode network (DMN) components. For each group and each pipeline, we extracted and z-transformed the seed BOLD time course and selected the top 10% time points with the highest activation. Consensus clustering revealed k=4 as the best-fitting k and by using k-means clustering spatial states of the four CAPs were then summarized in spatial z-maps and temporal occurrences (i.e. how many times a given CAP is expressed throughout the time course).

### 2.6 Statistical analysis

Within each group of participants, HC and PD, we evaluated separately whether static and dynamic resting-state networks (RSNs) were affected by the choice of head motion denoising strategy.

For the sFC analysis, we performed a dual regression to investigate differences in RSNs related to pre-processing pipeline in each group. To do so, based on our a-priori hypotheses, a paired t-test, randomized with permutation testing, was performed on 10 ICs with Bonferroni correction for multiple comparisons (10 ICs tested for 2 contrasts of interest: p-value< 0.0025) to detect network changes due to head motion controlling pipeline. Comparisons were done voxel-wise taking the standard *mc* as a reference pipeline and contrasting all the other pipeline to it (for HC: *mc>DVARS, mc>AROMA*; for PD: *mc>DVARS, mc>AROMA, mc>FIX*) in both contrasts’ directions.

For the transient-based dFC analysis, to understand voxels-wise differences between pipelines multiple paired t-tests were performed in randomise ^32^ across iCAPs of interest. Results are reported with Bonferroni correction considering dynamic network comparisons and 2 contrast of interest (respectively for HC group: [*AROMA* vs *mc, AROMA* vs *DVARS* and *mc* vs *DVARS* p_FWE_<0.00416] whereas for the PD group: *[AROMA* vs *mc, AROMA* vs *DVARS* and *FIX* vs *AROMA* p_FWE_<0.0125; *mc* vs *DVARS, FIX* vs *mc* and *FIX* vs *DVARS* p<_FWE_0.00625]; Figure S3). Furthermore, iCAPs durations between different pipelines were compared using a paired t-test for the two different runs described above for both groups (HC and PD separately, respectively contrasting *<u>mc</u>*<u>-</u>*<u>DVARS</u>* and *<u>mc</u>*<u>-</u>*<u>AROMA</u>* and *<u>mc</u>*<u>-FIX</u> for the PD only). The p-values were subsequently corrected for multiple comparisons with false discovery rate (FDR).

For the peak-based dFC analysis performed in the PD sample, CAP properties from different pipelines were compared by using a paired t-test for the two different runs described above (respectively contrasting *<u>mc</u>*<u>-</u>*<u>DVARS</u>* and *<u>mc</u>*<u>-</u>*<u>AROMA</u>* and *<u>mc</u>*<u>-FIX</u>) for occurrences, betweenness centrality, in degree, out degree and resilience and then correcting for multiple comparisons with FDR.

Finally, to gain insights into the effect of *AROMA* with respect to *mc*, a two-sample unpaired t-test was performed for both groups on the algebraic sum of the Independent Components labelled as noise and regressed out at single subject level by *AROMA*.

## 3. Results

### 3.1 Assessment of head motion

The amount of head-motion (both estimated as mean framewise displacement (FD) and DVARS) estimated during the resting state fMRI acquisitions was significantly higher in the PD group relative to the HC group: p_FD_<0.001, t-value_FD_=-7.5; p_DVARS_<0.001, t-value_DVARS_=-7.9 (Table 1; Figure S1).

**Table 1:** Head motion assessment of the sample.

| Population | DVARs<br>(mean±SD) | FD (mm)<br>(mean±SD) |
| --- | --- | --- |
| Healthy controls | 30±5 | 0.1±0.03 |
| Parkinson's disease | 45±7 | 0.3±1.2 |

### 3.2 Static functional connectivity

Despite the fact that the two groups showed significantly different natural head motion levels, the resting state networks (RSNs) derived from static FC approaches were rather robust as function to head motion denoising strategies (Figure 2). For the HC group (N=20): (i) the RSNs derived from *mc* and *DVARS* showed no significant differences; (ii) whereas relative to *mc, AROMA* shows significant voxel-wise differences for all RSNs but in brain regions expected to be motion-sensitive in both groups (Figure S2, Panel A). This can be interpreted as if *mc* denoising maintains a higher level of static FC in edge gray matter voxels relative to *AROMA* denoising. For what concerns PD (N=20): (i) both the comparisons between *mc* and *DVARS* and *mc* and *AROMA* shows significant voxel-wise differences (Figure S2, Panel B).; (ii) however RSNs derived from *mc* and *FIX* showed no significant differences. This can be interpreted as if *FIX* denoising maintains a within network static FC, comparable to the one seen using *mc*.

**Figure 2:**
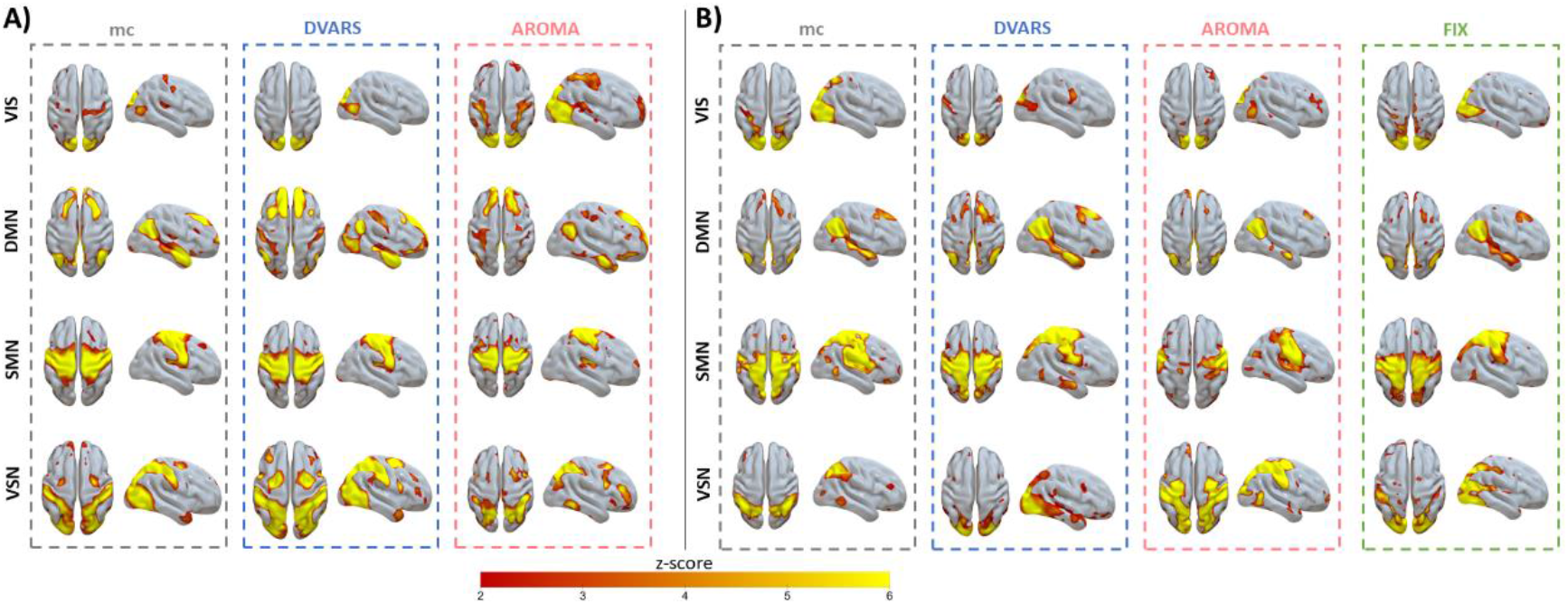
Head motion correction effects on static intrinsic functional networks in healthy controls and Parkinson’s disease. Spatial patterns of the independent components in the healthy control group (Panel A) and Parkinson’s disease (Panel B) were retrieved from all subjects in all sessions (gray: *mc*, blue: *DVARS*; red: *AROMA*, green: *FIX*).

### 3.3 Dynamic functional connectivity

#### 3.3.2 Transient-based analysis

Regarding dynamic FC in the HC group, from a spatial point of view, the three motion correction pipelines retrieved iCAPs maps of well-known resting-state networks (Figure 3; N=20; Table S1). However, while some iCAP networks appear to be robust as function as denoising strategy (auditory, visual, DMN, insula), other iCAPs are sensitive to denoising strategy (anterior Salience Network, aSN, and the visuospatial network, VSN), as can be qualitatively seen in Figure 3. In other words, even in a group of healthy volunteers with low head motion, the choice of head motion denoising strategy affects the spatial maps of transient-based dynamic networks.

**Figure 3:**
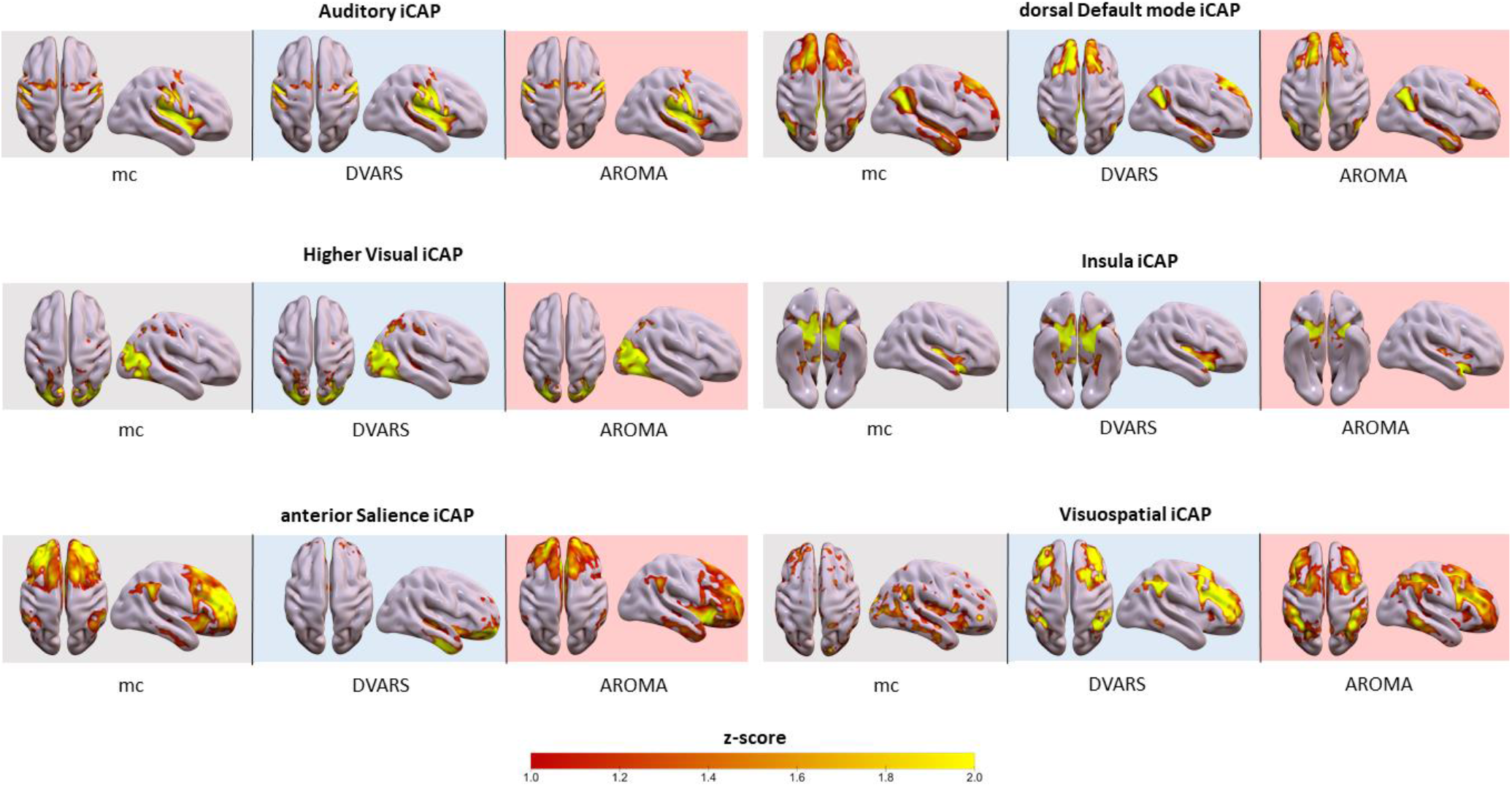
Head motion correction effects on dynamic intrinsic functional networks in healthy controls. Spatial patterns of the relevant innovation-driven coactivation patterns (iCAPs) in the healthy control group were retrieved from all subjects in all sessions (gray: *mc*, blue: *DVARS*; red: *AROMA*).

These effects of pipeline in the iCAPs of the HC group were further investigated by doing voxel-wise comparisons of the iCAP dynamic networks (Figure S3; Panel A), which showed: (i) lower connectivity in dorsal default mode network (dDMN) in precuneal regions in *DVARS* while compared to *mc* (p_FWE_< 0.01); (ii) lower connectivity in insula network in thalamic regions in *AROMA* while compared to *DVARS* (p_FWE_< 0.01); (iii) higher connectivity for the VSN network in both *AROMA* and *DVARS* compared to *mc*, in regions characteristic of the network itself, as demonstrated by the poor accuracy of the network detection (p_FWE_< 0.01); (iv) lower connectivity in frontal regions of the VSN while comparing *AROMA* to *mc* (p_FWE_< 0.01). All the reported results are family-wise errors (FWE)-corrected across voxels and Bonferroni corrected across different networks and contrast of interests.

When considering the temporal properties of the dynamic networks in the HC group, we found that the choice of head motion denoising strategy affects the percent duration of the various networks. We observed differences in engagement of a given state throughout the time course for the different runs (Figure 4): (i) for Auditory, Visual, aSN and dDMN total duration is significantly increased in *mc* compared to *DVARS* and compared to *AROMA* (p_FDR_<0.05), which also show a disruption in the Insula network, (ii) on the other hand, the VSN, which was poorly spatially retrieved with *mc*, had a reduced total duration compared to *DVARS* and *AROMA* (p_FDR_<0.05).

**Figure 4:**
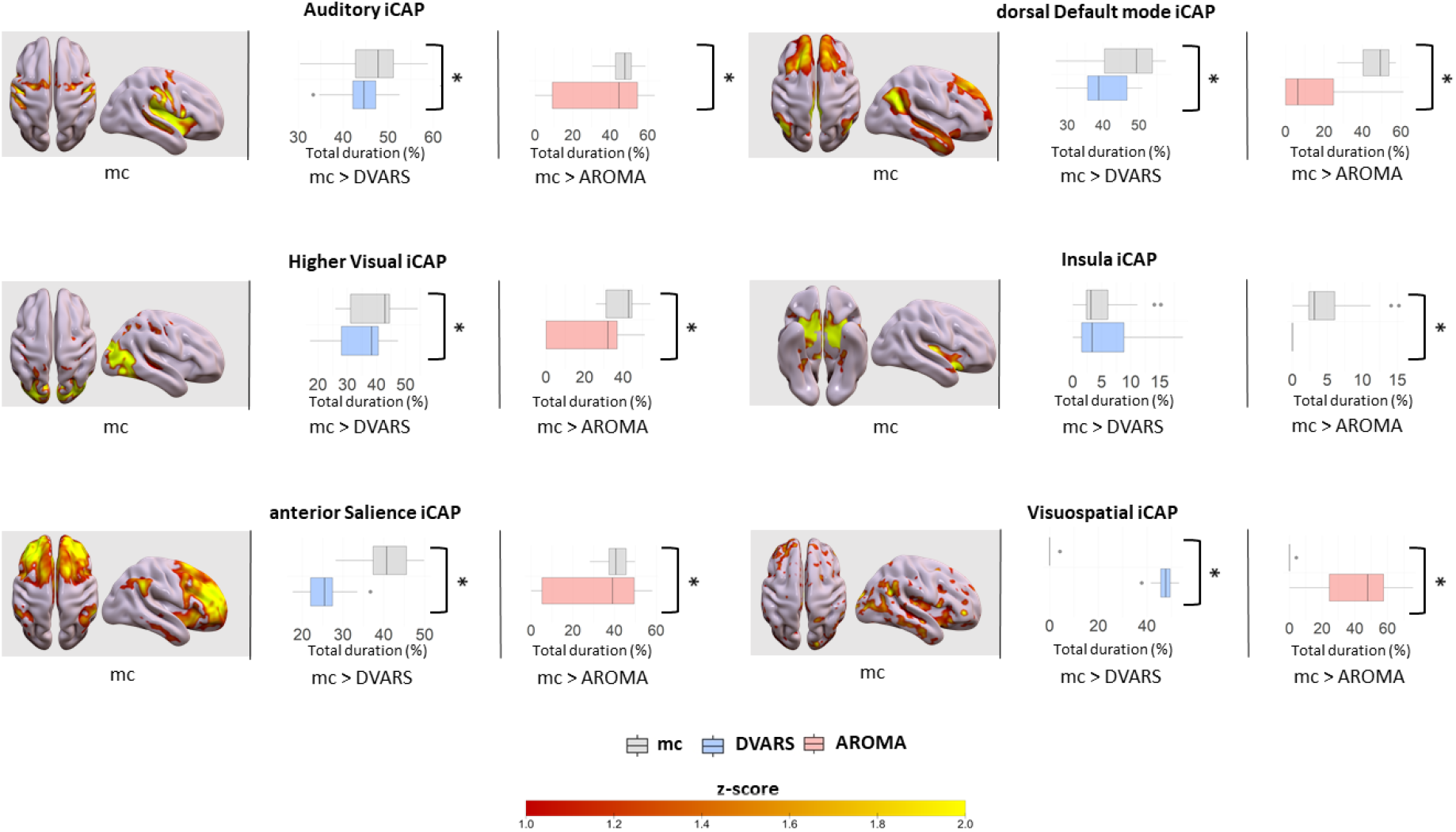
Head motion correction effects on the duration of intrinsic dynamic functional networks in healthy controls. For each iCAPs (spatial maps from *mc*), horizontal boxplots show the innovation frames per session in the healthy control group as a percentage of the total scanning time, highlighting the differences between: (i) *mc* vs *DVARS denoising*(*=p<0.05_FDR_) in the second column, (ii) *mc* vs *AROMA* denoising(*=p<0.05_FDR_) in the third column.

In the PD cohort, the spatial dynamic FC maps are severely affected by the motion correction pipeline (Figure 5; Table S1), resulting in a lower number of networks detected. Spatial distribution of brain networks is affected when choosing *AROMA* (two iCAPs detected) compared to *mc* and *DVARS* (four iCAPs detected). The *FIX* pipeline is the only denoising method enabling the detection of a higher number of consistent iCAPs compared to *mc* (five iCAPs detected). Therefore only 5 out of the 10 iCAPs (displayed in Figure 5), gave the resemblance of well-known resting state networks whereas the remaining iCAPs were characterized by noisy sparse activation.

**Figure 5:**
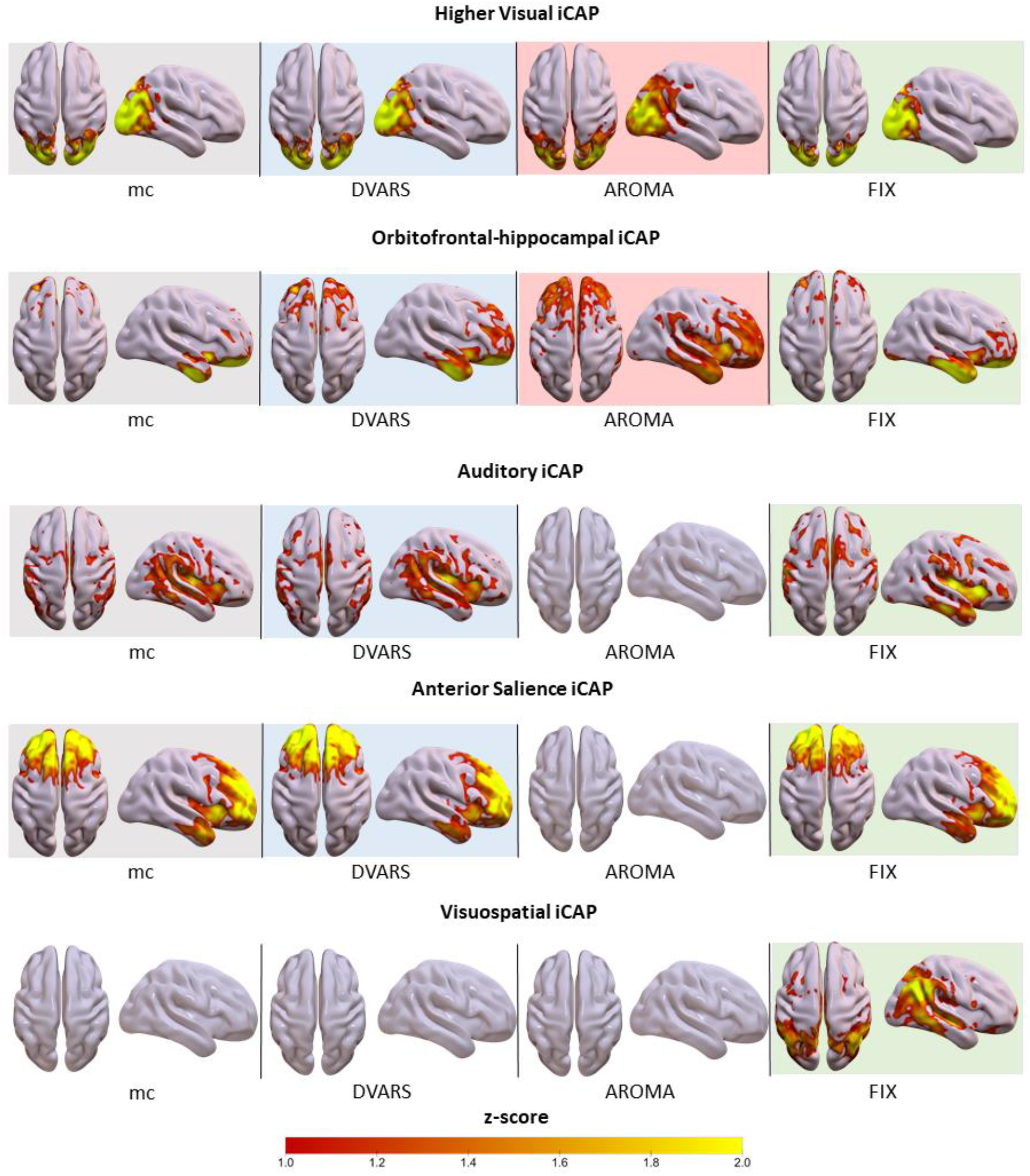
Head motion correction effects on the duration of intrinsic dynamic functional networks in Parkinson’s disease. Spatial patterns of the relevant innovation-driven coactivation patterns (iCAPs) in PD group retrieved from all subjects in different sessions (gray: *mc*, blue: *DVARS*; red: *AROMA*, green: *FIX*). Conditions in which no iCAPs were identified are left without maps.

Voxel-wise comparisons of dynamic networks (Figure S3; Panel B) revealed that: (i) lower connectivity in the Auditory network was found in cuneal and lateral regions while comparing *FIX* to both *mc* and *DVARS* (p_FWE_< 0.01); (ii) lower connectivity was also found in the Orbitofrontal-hippocampal network in orbitofrontal regions while comparing *AROMA* to both *mc* and *DVARS* (p_FWE_< 0.01), whereas higher connectivity was found within the same network in left temporal regions while comparing *FIX* to *AROMA* (p_FWE_< 0.01). Interestingly none of the detected iCAPs was significantly spatially different while comparing *DVARS* to *mc* (p_FWE_> 0.01). All the reported results are FWE-corrected across voxels and Bonferroni corrected across different networks and contrast of interests.

From a temporal perspective, dynamic FC networks in the PD group showed differences in engagement of a given state throughout the time course for the four different runs (Figure 6): (i) for Auditory network total duration is increased in mc compared to DVARS and *FIX* (p_FDR_<0.05), (ii) on the other hand, the Orbitofrontal-hippocampal had a reduced total duration in *mc* compared to *DVARS* and an increased total duration compared to *AROMA* (p_FDR_<0.05).

**Figure 6:**
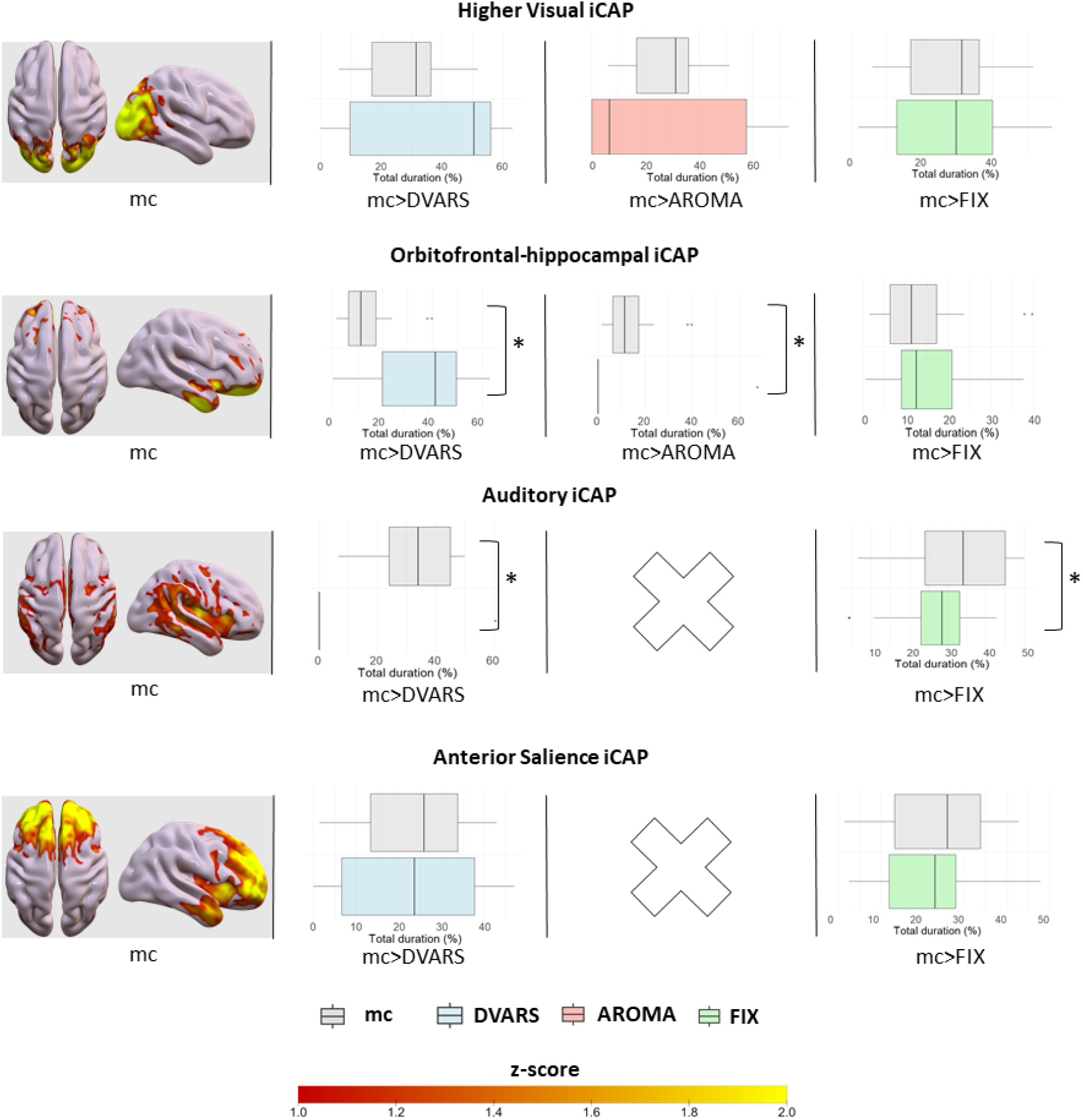
Head motion correction effects on the duration of intrinsic dynamic functional networks in Parkinson’s disease participants. For each iCAPs (spatial maps from *mc* denoising), horizontal boxplots show the innovation frames per session in the Parkinson’s disease group as a percentage of the total scanning time, highlighting the differences between: (i) *mc* vs *DVARS denoising* (*=p<0.05_FDR_) in the second column, (ii) *mc* vs *AROMA denoising* (*=p<0.05_FDR_) in the third column, (iii) *mc* vs *FIX denoising* (*=p<0.05_FDR_) in the fourth column. No AROMA iCAPs were detected for the Auditory and Anterior salience networks.

Voxel-wise statistical maps of the noise components removed by *AROMA* (Figure S4) show a clear increase in the effect of *AROMA* denoising in PD compared to HC (p_FWE_<0.05).

In summary, transient-based dynamic intrinsic functional connectivity analysis is sensitive to the choice of head motion denoising strategy. In a population of young healthy subjects with low amounts of head motion, the pipeline choice affects some of the spatial iCAPs and the percent duration of the various networks. In a population of PD participants, instead, the effects are rather more drastic, and it becomes challenging to actually estimate iCAPs.

#### 3.3.2 Peak-based analysis

The transient-based iCAPs results in the PD group showed that it was not possible to detect the expected dynamic networks present in the HC group, such as the DMN. We hypothesized that a potential reason for this may be related to the HRF assumption that is built into the iCAPs method for its temporal deconvolution step. In addition, functional MRI activity transients are detected using the derivative of this signal, which can further amplify the effect of noise. We tested this on the DMN network by using the PCC seed-based CAP technique for dynamic connectivity in the PD group. Using this, and irrespective of head denoising strategy, we were able to detect comparable dynamic DMN co-activating networks.

Voxelwise cross-correlation between different DMN CAPs z-scored maps revealed high anatomical concordance across pipelines in PD cohort (Figure 7) (*mc* vs *DVARS*: r-value=0.99; *mc* vs *AROMA*: r-value=0.90; *mc* vs *FIX*: r-value=0.97).

**Figure 7:**
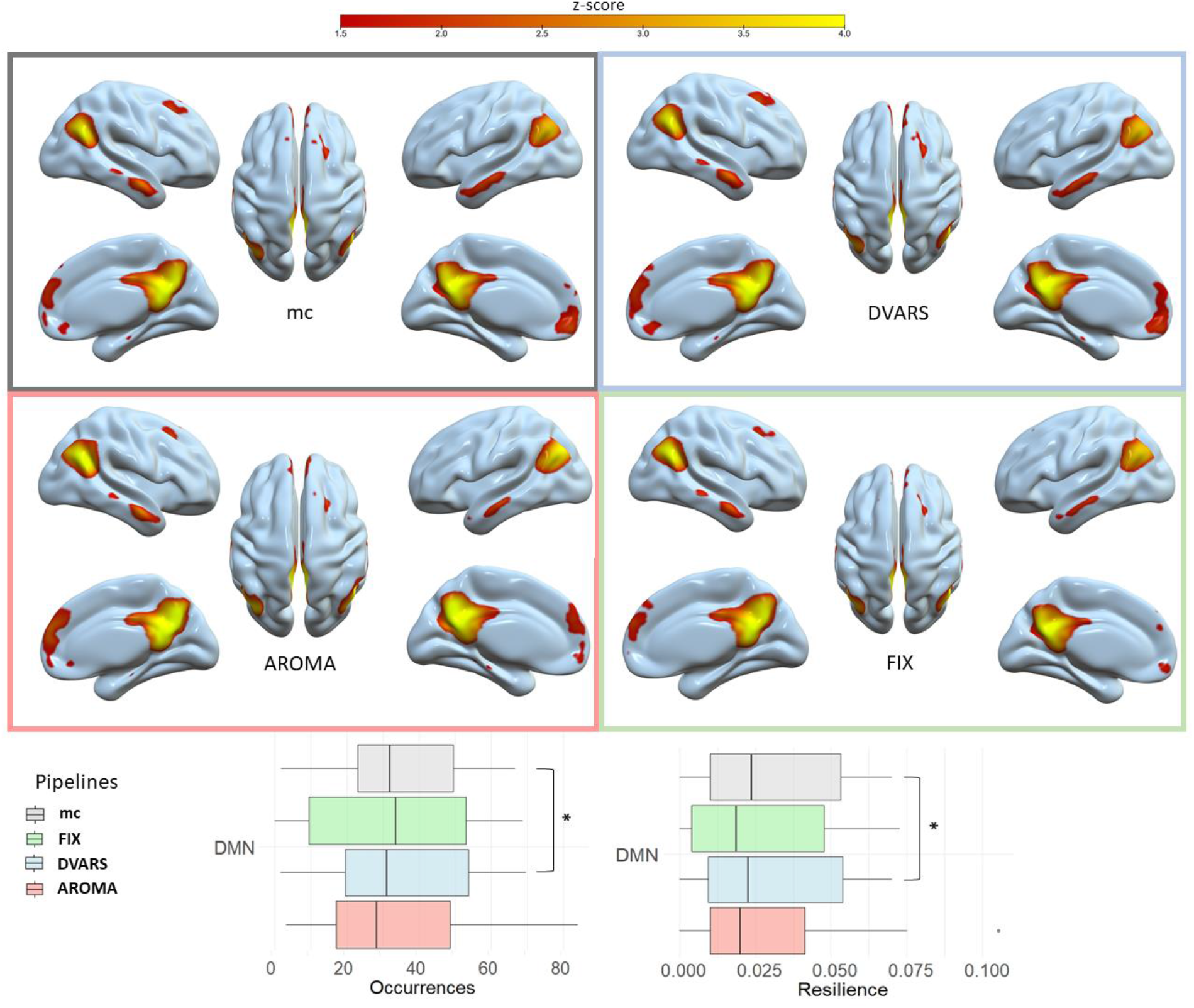
Intrinsic default mode network (DMN) co-activation patterns (CAPs) in Parkinson’s disease participants across pre-processing pipelines. Surface display of DMN CAPs in, starting from upper left: *mc* (gray); *DVARS* (light blue); *AROMA* (light pink) and *FIX* (light green). Boxplots display respectively occurrences and resilience of the DMN CAP (p-value_FDR_<0.05).

Moreover, by considering the occurrence of the DMN across subjects, *FIX* resulted to be the head motion denoising pipeline with higher z-score median and low z-score standard deviation across voxels compared to other pipelines, showing higher temporal stability in the detection of the network.

In terms of DMN CAP temporal properties (Table S2), we found no significant differences when comparing the results derived from using the *mc* to either of the *AROMA* or *FIX* denoising pipelines. Instead, significant differences were found when comparing temporal properties from *mc* relative to *DVARS* (Table S4): (i) a decrease in DMN occurrences with *mc* (i.e. how many times the CAP is expressed across the time course, p-value_FDR_<0.05, t-value=-3.5) and a decreased in DMN resilience with *mc* (i.e. a stability index expressing the likelihood to remain in the same CAP configuration across time, p-value_FDR_<0.05, t-value=-3.1).

## 4. Discussion

In this study, within two groups of subjects having different head motion levels (HC and PD), we evaluated how standard *mc, DVARS, AROMA* and *FIX* denoising methods differentially affect intrinsic static and dynamic FC metrics.

We replicated previous results in the field, showing that large head motion and denoising strategies do affect intrinsic FC estimates ^6,33^. Until now, such evaluations have been done mostly applying static FC estimations. In this study, we extend those results to the consideration of framewise dynamic FC strategies, both transient-and peak-based. Our main finding is that both of these dynamic FC methods are affected by both the level of head motion and the retrospective denoising methods used to mitigate motion effects. Indeed, we demonstrate that even in a group of HC with low level of head motion, dynamic FC results can be affected by denoising pipelines. Whereas, if the group shows larger head motion such as in our PD group, dynamic FC can completely fail to capture transients-based networks. On the other hand, the use of peak-based dynamic FC can help recovering networks in the PD group, but these can still show temporal features of dynamic networks that are dependent on head motion denoising pipeline.

### 4.1 Head motion correction pipelines: effects on the estimation of intrinsic functional connectivity metrics

When looking at the effect of head motion correction in the two samples (HC and PD), it is clear that different denoising strategies will distinctively affect the results derived downstream.

Previous studies^10^ suggested that volume censoring (i.e. “scrubbing”) could be beneficial in the context of dynamic FC, a viable approach that implicitly underestimates the temporal relationship of timepoints ^11^. Indeed, in this study, we demonstrate that volume de-weighting alone (*DVARS*) is sufficient to alter temporal properties of iCAPs (e.g., total network duration) and CAPs (e.g., network occurrences and resilience), thus extending previous reports on sliding-window approaches ^34,35^. Moreover, we showed that in a high-motion population even a subtle scrubbing approach such as DVARS can cause a decrease in within network static FC.

On the other hand, data-driven decomposition approaches for denoising such as *AROMA* and *FIX*, may be more specific in filtering motion related temporal dynamics of brain MR signals ^36,37^. For what concerns static FC, we found differences in within-network FC in both populations while comparing *mc* with *AROMA*, with the PD cohort exhibiting larger changes in key network nodes. Whereas when comparing *mc* with *FIX* in the PD sample, no significant differences were found.

In fact, in the PD group, where the denoising was performed separately for *AROMA* and *FIX*, we found that *AROMA* removed a lot more “noise” and probably as a consequence detected fewer dynamic networks. It is possible that *AROMA* performed in a suboptimal way on our PD dataset (TR=1s) because the detection and elimination of head motion components has been originally developed and validated under different experimental conditions (TR= 1.96s^15^), which differentially affects the spectral distribution of noise components. The higher sampling rate of our experiment most likely allows us to maintain low frequency BOLD components that have less aliasing from higher frequency components, like from head motion. Therefore, the use of *AROMA* may remove some low frequency components that are wrongly interpreted as noise ^38,39^.

Instead, in the case of *FIX*, which in PD enables the detection of a higher number of iCAPs compared to *mc* by preserving their temporal properties, ICA-denoising seems to be more effective and not disruptive. In this study, the *FIX* classifier, later tested on PD, was trained on manually labelled noisy components detected by four experts in our HC sample. This enabled the definition of components resembling “noise”, being it physiological or related to head motion, specific to our acquisition protocol. Importantly, the acquisition protocol was identical in the training (HC) and testing (PD) sample.

### 4.2 Transient-based dynamic connectivity is sensitive to head motion and head motion denoising pipeline

We showed for the first time that frame-wise dynamic FC is sensitive to both the level of head motion in the group and the choice of head motion denoising pipeline.

Our results demonstrated how even in the case of small-head motion (FD <0.1 mm) frame-wise techniques, especially iCAPs, are susceptible to different head motion correction. Indeed, in HC standard head motion correction (*mc*) was not sufficient to dynamically detect networks such as the VSN. However, more aggressive denoising techniques (i.e. *AROMA*) resulted in changes of temporal properties of several iCAPs (such as temporal durations) which are closer to zero (i.e. absent) for a large part of the sample even if characterized by a low motion level, as previously stated ^40^. Besides, if we look at populations with inherited large motion levels, implications are even worse. In the PD sample, iCAPs are spatially disturbed with simple outliers’ volume de-weighting (*DVARS*) whereas large parts of networks are absent for more disruptive denoising such as *AROMA*. Therefore, it seems that in the presence of a higher proportion of corrupted functional volumes, two factors may contribute to the loss of sensitivity to detect dynamic networks. On one hand, the removal of noisy components from the original signal interferes with the proper reconstruction of spatial and temporal properties of iCAPs. On the other hand, the simple regression of volumes from the time-course disables the capability of the algorithm to detect these fluctuations. On the contrary, applying a denoising pipeline that is more specific to the sample (i.e. FIX), appears to help preserve the signal temporal variance and frequency content, thus improving the estimation of the iCAPs ^41^ by boosting network identifiability.

### 4.3 The use of co-activation patterns enables the detection of well-known functional networks even in high motion populations

We found that the iCAPs algorithm was not able to detect the dynamics of well-known RSNs, such as the DMN, in samples characterized by large head motion (PD). Thinking that this may be related to problems with the HRF deconvolution in a noisy time series, we explored the use of an alternative framewise dynamic method. We applied the CAPs technique which rather than looking for transient framewise fluctuations, it attempts to detect peaks in the single frame. We found that by selecting a precise region of interest, even in the presence of large head motion, the CAPs framework is capable of detecting DMN co-activation and deactivation over the time course. However, we showed that the temporal properties of the DMN are still differentially affected by the choice of denoising pipeline, especially when using *DVARS*. This may be partially explained by the fact that CAPs correlations are largely driven by the spatial nonspecific correlation components of global signals ^42^. The close relationship between spatially non-specific CAPs and head motion can be strongly affected by the denoising pipeline. In fact, we observed that the pipeline affects the classification of single frame peaks over the time course as neurally relevant or not CAPs. Indeed, *DVARS* head motion correction is thought to be also sensitive to global signal changes in the BOLD content not related to motion ^5^, resulting in considerable effects on temporal CAPs compared to other denoising techniques.

Moreover, it is also relevant to keep in mind that iCAPs and CAPs, being framewise dynamic techniques, are largely affected by changes in transient arousal level of the subjects, which are usually accompanied by head motion peaks ^34^.

### 4.4 Limitations

This study has several limitations. We did not aim to directly compare intrinsic functional activity across the HC and PD groups, which were not age matched. Instead, the focus was on quantifying how static and dynamic connectivity networks were affected by head motion denoising pipeline as an independent variable. Future studies are needed to generalize our results to other age ranges of our groups. Also, in terms of head motion denoising methods, our goal was not to explore all possible head motion correction methods but rather to investigate if some of the most used techniques differentially affect FC representations, especially with dynamic connectivity approaches. Using the healthy young volunteer’s data as a training set for *FIX* denoising of the Parkinson’s group may be suboptimal because of the age differences across groups. The peak-based dynamic connectivity was limited to the PD group only and considering only one seed, the posterior cingulate cortex, in order to evaluate the possibility of detecting the default mode network. The healthy group was not evaluated with the peak-based dynamic method because the known networks were retrieved with the transient-based approach. Further, our estimation of TVC patterns was limited to frame-wise approaches. Further studies are needed to better understand the effects of head motion denoising in sliding-window correlation and other dynamic algorithms ^28,43^.

## 5. Conclusions

To the best of our knowledge this is the first study comparing how head motion denoising strategies affects static and dynamic functional intrinsic BOLD connectivity in two populations with very different natural head motion properties. We found that static intrinsic FC metrics are mostly robust across motion denoising strategies in both young healthy controls and Parkinson’s participants. This is consistent with and supports previous studies ^44,45^. However, dynamic FC metrics were very sensitive to denoising methods, in both groups but critically in the PD group. Therefore, further work is needed to better understand: (i) how to retain useful information from a BOLD time-series in high motion populations, (ii) and how to disentangle subject-specific dynamic head motion signatures potentially retaining relevant neural information ^13,41^. Ultimately, our study contributes to emphasizing the importance of head motion in the context of novel dynamic FC studies. To facilitate the comparison and reproducibility of such studies it remains crucial to report both the head motion properties of the populations involved and the details of the denoising strategies used ^46^.

## Acknowledgments

This study was supported by the ISMRM Exchange Award 2021-2022 “*Investigating in-vivo human brain dynamic connectivity with fast fMRI*”, the Dipartimento di Eccellenza project 2018-2022 (Italian Ministry of Education, University and Research), and the Caritro foundation project “Tango, una terapia complementare per la malattia di Parkinson”.

