## Supplementary materials for "Head motion correction shapes functional network estimates: evidence from healthy and Parkinson’s disease cohorts"

#### Supplementary figures

**<u>Figure S1</u>. Head motion levels.** Panel <u>A</u> displays mean framewise displacement (FD, mm) across volumes. Panel <u>B</u> displays boxplots of framewise displacement at the single subject level in healthy controls (HC). Panel <u>C</u> displays boxplots of framewise displacement at the single subject level in Parkinson's disease patients (PD).

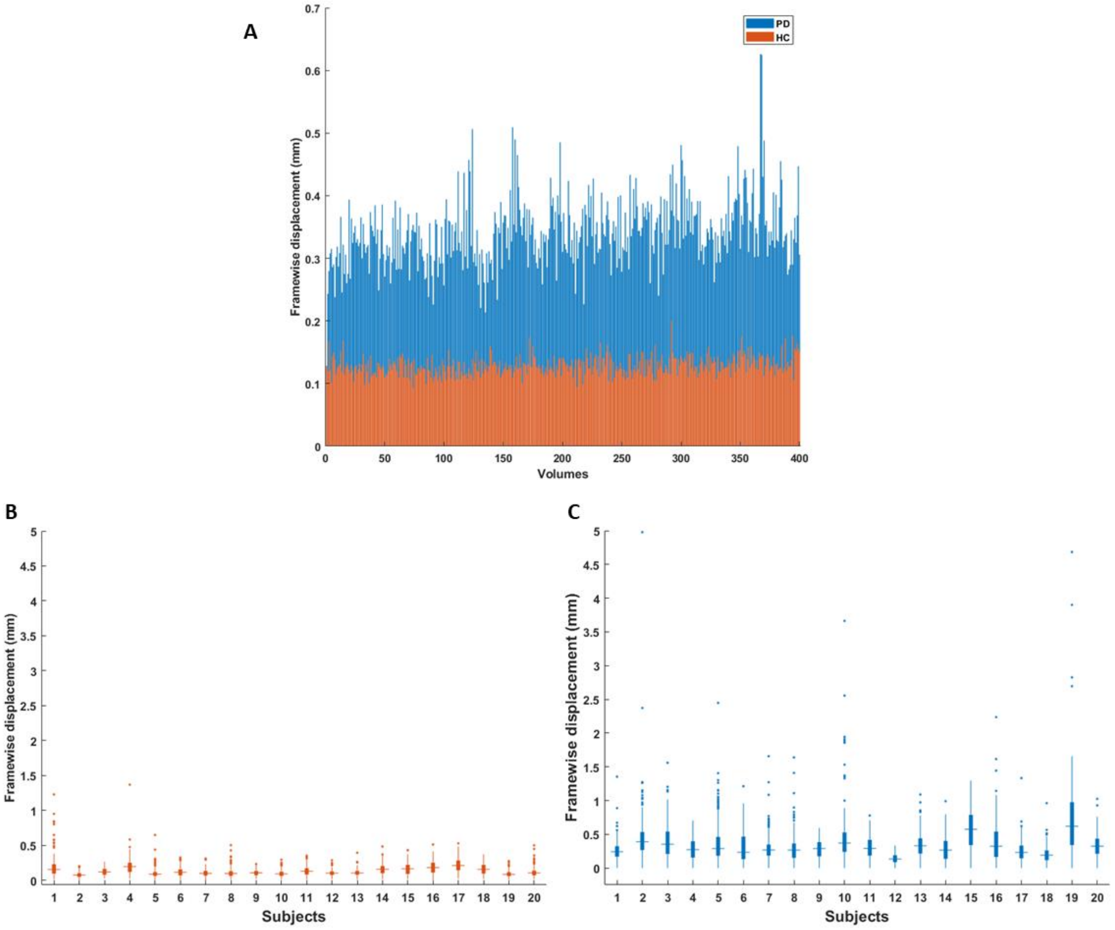

**<u>Figure S2:</u> Head motion correction effects on static intrinsic functional connectivity (sFC) networks in healthy controls (HC) and Parkinson's disease (PD) groups.** Panels A and B show, for HC and PD respectively, the spatial patterns of voxels where *mc* derived resting state networks significantly differ ($p<0.001_{Bonferroni}$) relative to sFC networks derived the other denoising strategies (red-yellow maps where *mc* gives higher sFC, blue-light blue maps where *mc* gives lower sFC).

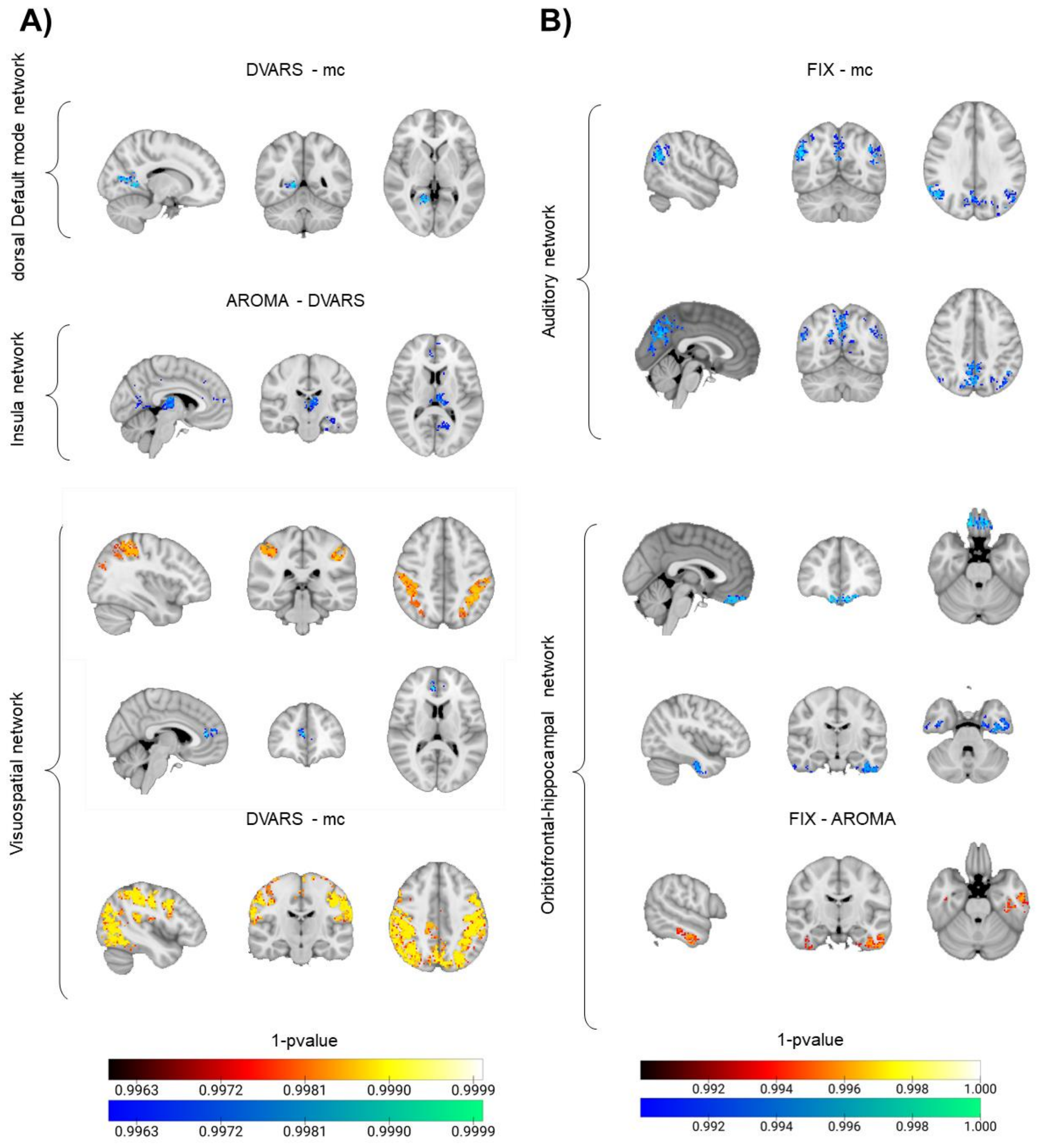


Abbreviations: VISUAL (Visual network), DMN (Default mode network), SMN (Somatosensory network), FPN (Fronto-parietal network), VSN (Visuospatial network).

**<u>Figure S3</u>: Voxel-wise effects of denoising pipelines in iCAPs maps.** Panel <u>A)</u> on the left-hand side displays voxel-wise differences in spatial localization of iCAPs for the tested contrasts [*DVARS-mc*; *AROMA-DVARS*; *AROMA-mc*]. Panel <u>B</u> on the right-hand side displays voxel-wise differences in spatial localization of iCAPs for the tested contrasts [*DVARS-mc*; *AROMA-DVARS*; *AROMA-mc*: *AROMA-FIX*; *FIX-DVARS: FIX-mc*].

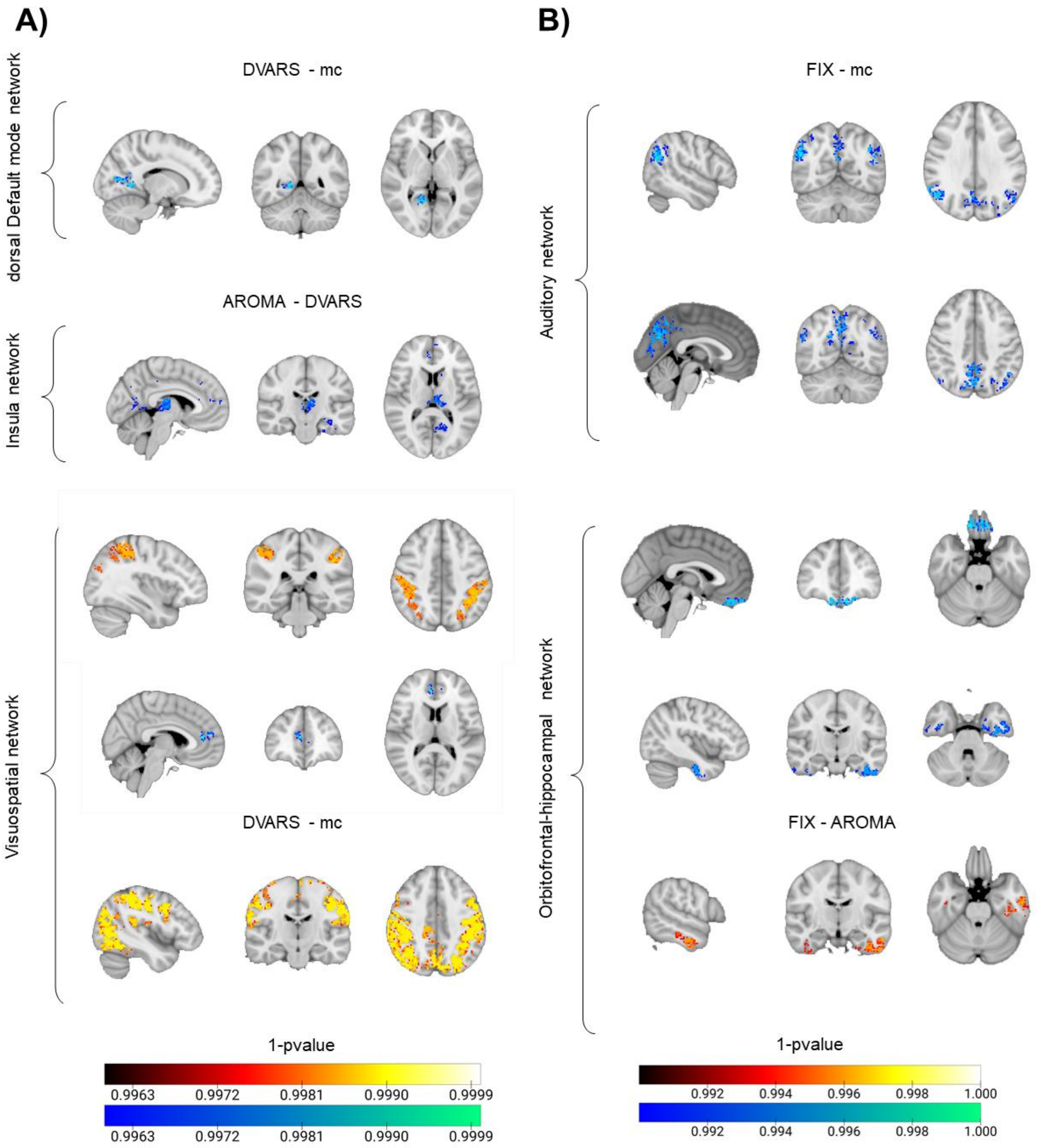

Figure S4: **Examination of removed noise components from intrinsic functional fMRI time series.** Connectivity component maps removed by *AROMA* pipelines within the healthy control (HC), Parkinson's disease (PD) groups, and their relative comparison.

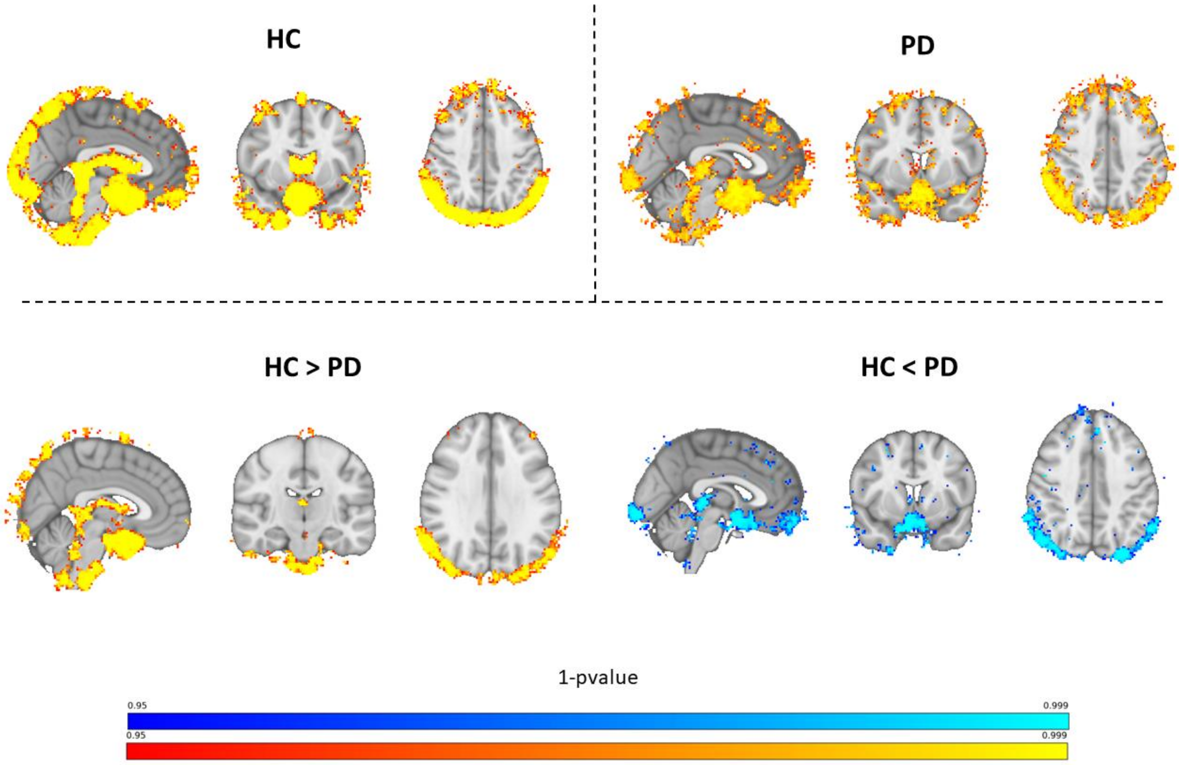

### Supplementary tables

**<u>Table S1</u>**. Voxel-wise cross-correlation values between iCAPs detected networks and well-known static resting-state networks from Shirer et al., 2012 in Healthy controls (HC) and Parkinson's disease patients (PD). The detected iCAPs of Insula (in HC) and Orbitofrontal-hippocampal networks (in PD) are not reported given the absence of corresponding static resting-state network.

| iCAP Network | Dataset | Pipeline | r-value |
|---|---|---|---|
| <u>Auditory</u> | HC | *mc* | 0.20 |
| | | *DVARS* | 0.20 |
| | | *AROMA* | 0.30 |
| | PD | *mc* | 0.12 |
| | | *DVARS* | 0.12 |
| | | *AROMA* | *N.A.* |
| | | *FIX* | 0.17 |
| <u>Higher Visual</u> | HC | *mc* | 0.22 |
| | | *DVARS* | 0.30 |
| | | *AROMA* | 0.5 |
| | PD | *mc* | 0.28 |
| | | *DVARS* | 0.27 |
| | | *AROMA* | 0.20 |
| | | *FIX* | 0.32 |

| | | | |
|---|---|---|---|
| anterior Salience | HC | *mc* | 0.32 |
| | | *DVARS* | 0.12 |
| | | *AROMA* | 0.20 |
| | PD | *mc* | 0.24 |
| | | *DVARS* | 0.24 |
| | | *AROMA* | *N.A.* |
| | | *FIX* | 0.22 |
| dorsal Default mode | HC | *mc* | 0.44 |
| | | *DVARS* | 0.40 |
| | | *AROMA* | 0.42 |
| | PD | *mc* | *N.A.* |
| | | *DVARS* | *N.A.* |
| | | *AROMA* | *N.A.* |
| | | *FIX* | *N.A.* |
| Visuospatial | HC | *mc* | 0.20 |
| | | *DVARS* | 0.24 |
| | | *AROMA* | 0.33 |
| | PD | *mc* | *N.A.* |
| | | *DVARS* | *N.A.* |
| | | *AROMA* | *N.A.* |
| | | *FIX* | 0.14 |

**<u>Table S2.</u>** Summarizing stats of comparison between temporal properties of DMN CAPs across pipelines in the Parkinson's patient's cohort (*=p-value$_{FDR}$<0.05, bold=p-value<0.05).

| Comparison | Metric | p-value | t-value |
|---|---|---|---|
| *mc>DVARS* | Occurrences | **0.0024*** | -3.4940 |
| | Betweenness Centrality | 0.2617 | 1.1567 |
| | Resilience | **0.0058*** | -3.1066 |
| | In degree | 0.6058 | -0.5247 |
| | Out degree | 1 | 0 |
| *mc>AROMA* | Occurrences | 0.6626 | -0.4432 |
| | Betweenness Centrality | 0.8884 | 0.1422 |
| | Resilience | 0.4991 | -0.6890 |
| | In degree | 0.2883 | 1.0924 |
| | Out degree | 0.2345 | 1.2280 |
| *mc>FIX* | Occurrences | 0.9432 | 0.0722 |
| | Betweenness Centrality | 0.3195 | 1.0222 |
| | Resilience | 0.3209 | -1.0193 |
| | In degree | **0.0217** | **2.5015** |
| | Out degree | 0.0828 | 1.8311 |
